# An efficient iDNA method for surveying rainforest mammals with carrion flies

**DOI:** 10.1101/2025.08.25.672224

**Authors:** Torrey W. Rodgers, Arianna Basto, Raider Castro, Erin Marcela Rivera Groves, Charles C. Y. Xu, Elena Hernández, Karen E. Mock, Adrian Forsyth

## Abstract

Metabarcoding of vertebrate DNA derived from invertebrates (iDNA) is a promising method for monitoring of mammal populations However, optimization of field and laboratory methods for efficient and economical sampling and analysis is needed. We used metabarcoding of carrion flies to survey for mammals at Los Amigos Biological Station in the Peruvian Amazon. We used an extraction-free, direct PCR approach to metabarcode vertebrate DNA, whereby buffer from tubes that contained flies for 3-5 hours was used directly in PCR, negating the need for labor-intensive DNA extraction. We also examined two different pooling methods to investigate the efficacy of pooling multiple fly samples to economize sequencing of large numbers of samples. Metabarcoding detected 27 mammalian taxa from 9 orders and 18 families plus four anuran taxa from three families from just 240 carrion flies collected over 3 days. Pooling of buffer from multiple flies prior to PCR resulted in the loss of many species compared to sequencing of individual flies. However, pooling of PCR product after an initial PCR from individual flies, but prior to indexing, resulted in species detection nearly identical to sequencing of individual flies. These methodological findings will contribute to the feasibility of iDNA metabarcoding for vertebrate monitoring and biodiversity conservation.

## Introduction

Environmental DNA (eDNA) is a powerful tool for biodiversity monitoring (Bohmann et al., 2014; Ruppert et al., 2019). Historically, vertebrate eDNA research focused primarily on aquatic species, however, eDNA methods have more recently been adapted for the study of terrestrial vertebrates (Newton et al., 2025). In terrestrial systems, invertebrate-derived DNA (iDNA) methods using invertebrates such as carrion flies (Calvignac-Spencer, Merkel, et al., 2013; Rodgers et al., 2017), dung beetles (Drinkwater et al., 2021; Saranholi et al., 2024) and leeches (Ji et al., 2022) allow for passive surveillance of terrestrial vertebrate communities. By acquiring traces of vertebrate DNA through interactions with feces, carcasses, blood or dead skin from live animals, these invertebrates can assist in the monitoring of terrestrial vertebrate assemblages. This methodology allows detection of vertebrate species presence without direct sightings or captures and offers an effective and non-invasive approach for surveys in tropical rainforest ecosystems (Rodgers et al., 2017) where high biodiversity, cryptic species, remote landscapes, and a rich canopy fauna may limit the efficacy of camera traps or other survey methods.

Although iDNA metabarcoding has been used effectively for vertebrate sampling (Calvignac-Spencer, Leendertz, et al., 2013; Drinkwater et al., 2021; Ji et al., 2022; T. W. Rodgers et al., 2017; Saranholi et al., 2024; Schubert et al., 2015), methodological advancements to increase efficiency are needed. DNA extraction is often one of the most time-consuming steps in eDNA or iDNA metabarcoding laboratory pipelines. For carrion fly metabarcoding, past methods required grinding whole flies into a slurry for DNA extraction (Calvignac-Spencer, Merkel, et al., 2013; T. W. Rodgers et al., 2017) or dissecting fly digestive tracts (Lee et al., 2016), processes that demand substantial effort. This study utilized carrion flies to survey mammalian diversity in the Amazon rainforest of Los Amigos Biological Station, Madre de Dios, Peru through iDNA metabarcoding. We tested a field method of netting carrion flies directly off a bag of pork bait to avoid having to catch flies in traps. We also tested an extraction-free, direct PCR approach for next generation sequencing of mammal DNA derived from carrion flies (Srivathsan et al., 2023), a method which reduced laboratory processing time by tens of hours by avoiding grinding, dissection, and DNA extraction altogether. Additionally, we compared two different methodologies for pooling fly samples during library preparation to determine the most efficient approach for large-scale biodiversity assessments where sequencing of individual flies is cost prohibitive.

## Methods

### Study site

Los Amigos Biological Station (LABS; also known as CICRA in previous literature), is located at 12°33’40.0” S, 70°05’46.4” W in the Amazon Basin at an elevational range from 225 to 296 meters above sea level in the Madre de Dios region of southeastern Peru (Figure 1). The sampling area was at the convergence of the Los Amigos and Madre de Dios rivers. This area includes a range of habitats including mature old growth forest on terra firme terraces of ancient, weathered soils of low pH, as well as both mature and successional forest found on the more fertile alluvial floodplains that are annually renewed by episodic flooding, where there are dense stands of *Guadua* bamboo and *Mauritia flexosa* palm swamps. Soils are primarily ultisols and inceptisols. The region has a tropical humid climate with a rainy season from November to February and a dry season from June to September. Annual rainfall ranges between 2700 and 3000 mm, and the mean temperature ranges from 21°C to 26 °C. The region supports an intact mammalian community that is currently unhunted and is typical of intact lowland rainforests in the western Amazon.

**Figure 1.**
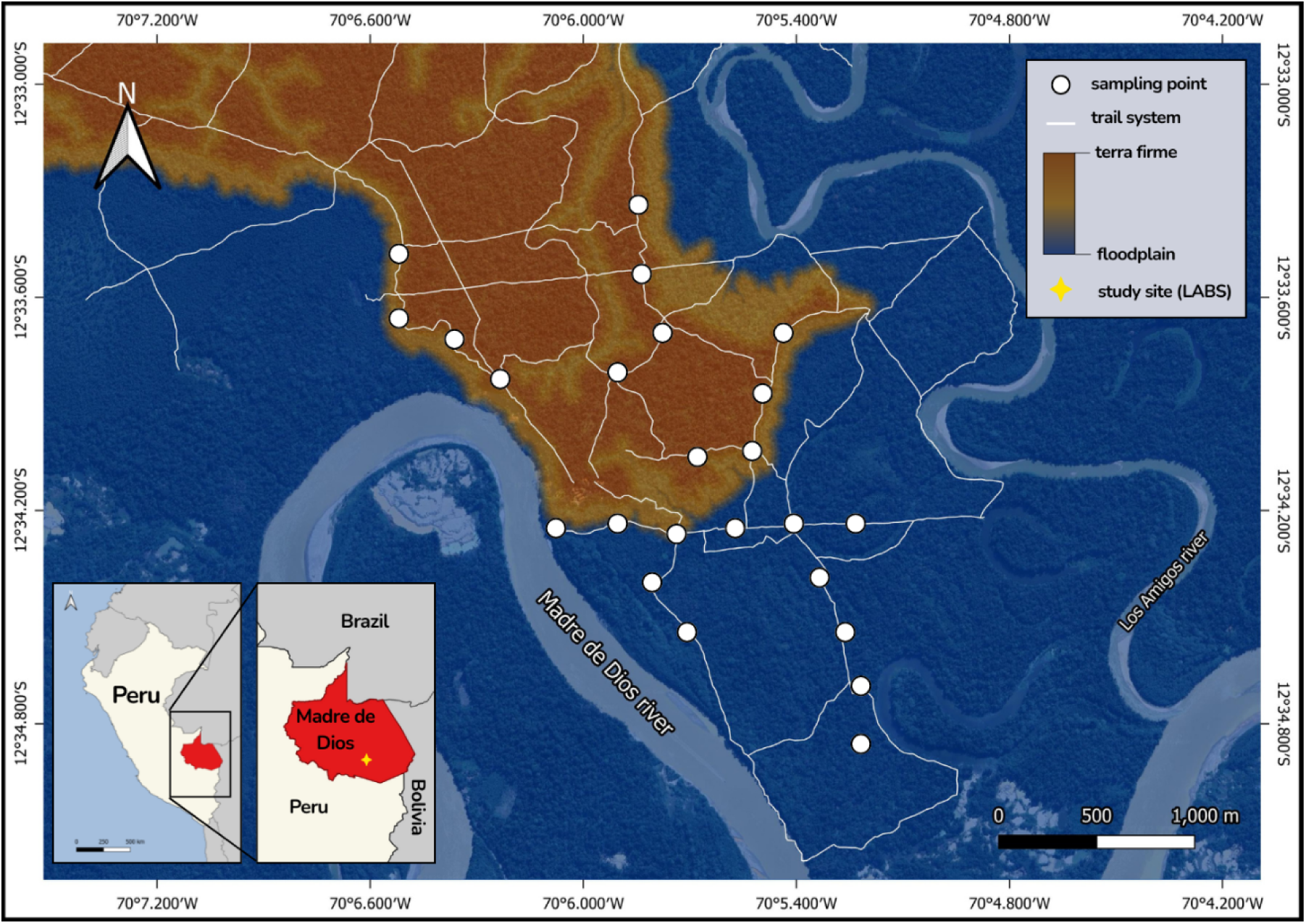
Map of study site for collection of carrion flies for iDNA metabarcoding to detect rainforest mammals at Los Amigos Biological Station, Madre de Dios, Peru.

### Carrion fly collection

Carrion flies were collected by netting directly off a white plastic grocery bag, tied loosely at the top, with an open Ziplock bag filled with rotting pork inside. Pork was rotted under the sun for at least 24 hours prior to sampling. Fly sampling was conducted on Nov 9-11, 2023. At each site, 10 flies were netted and individually placed directly into 2 ml or 1.5 ml sterile snap-cap microcentrifuge tubes. After 10 flies were collected from one site, a new site was selected 300 meters further along the trail. This was repeated four times each morning and afternoon for a total of 240 flies collected across 24 locations from six trail segments, with three segments in floodplain habitat and three in terra firma habitat (Fig. 1).

Flies were brought to the field laboratory and left in tubes at ambient temperature for 3-5 hours, after which they were removed from tubes and placed in EtOH for morphological species identification. Next, 4-5 blue-indicating silica gel beads were placed in each empty tube to dry the interior, and tubes were then placed in a −20 °C freezer until transport to the molecular laboratory. Prior to transport, silica gel beads were removed and disposed of, and dry, empty tubes were transported to the molecular ecology laboratory at Utah State University, where they were placed in a −70 °C freezer (Fig. 2A).

**Figure 2.**
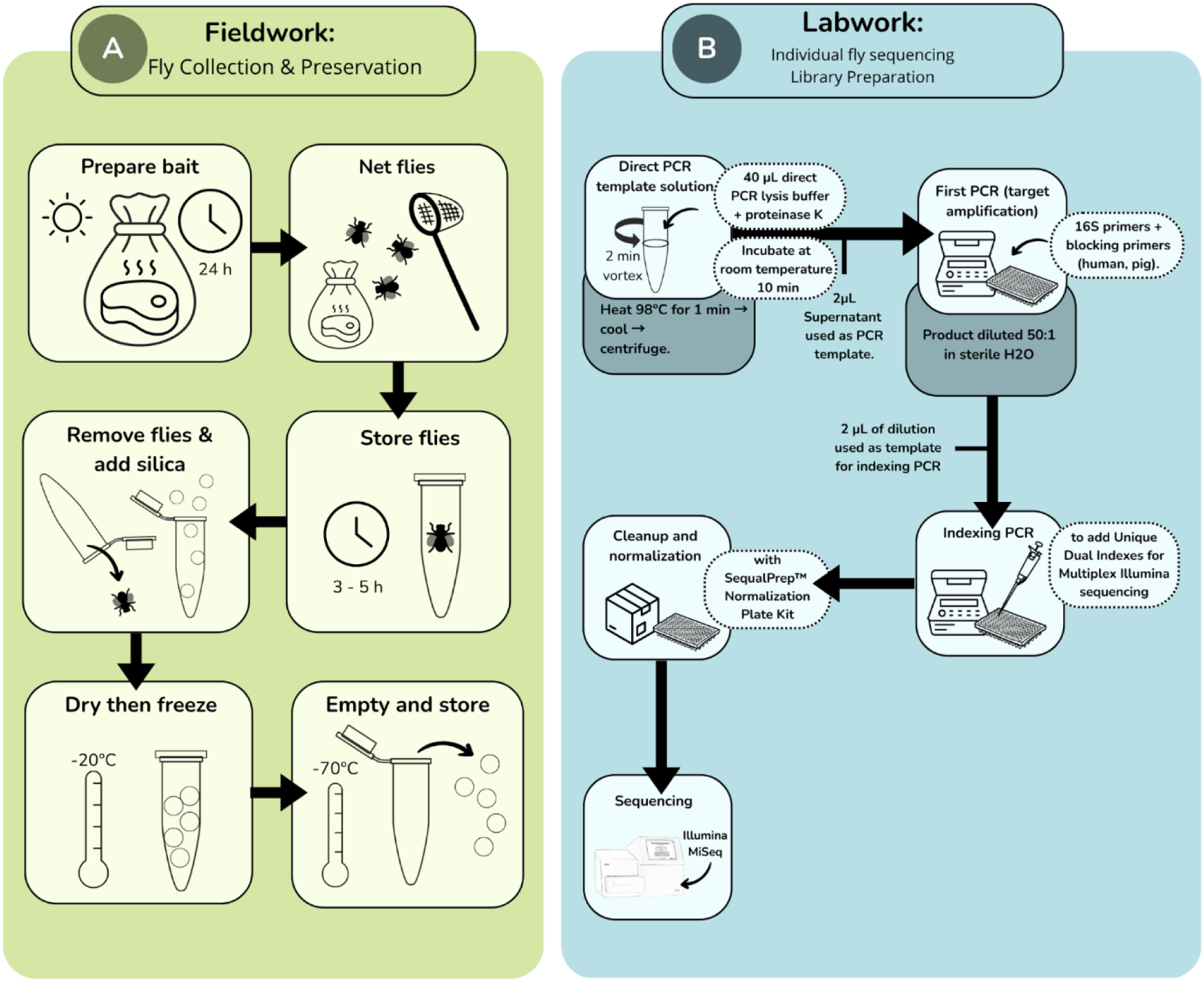
Workflow for a) carrion fly collection field work and B) laboratory work for sequencing of individual carrion flies for detection of rainforest mammals with iDNA from Los Amigos Biological Station, Madre de Dios, Peru.

### Laboratory methods

We used an extraction-free, direct PCR approach for DNA amplification (Fig. 2B). Forty microliters of direct PCR lysis buffer (supplied with Platinum™ Direct PCR Universal Master Mix, Invitrogen, Carlsbad, CA) with 1.2 μL of proteinase K were added to each empty microcentrifuge tube from fly sampling. Tubes were vortexed for 2 minutes, incubated at room temperature for 10 minutes, and then incubated on a heat block at 98 °C for one minute, before cooling to room temperature (as per manufacturers’ lysis protocol). Samples were then centrifuged for one minute, and the buffer template solution was added directly to PCR reactions.

PCR reactions contained 10 μL Platinum™ Direct PCR Universal Master Mix (Invitrogen, Carlsbad, CA), 4 μL Platinum™ GC Enhancer, 0.2 μM each of forward and reverse primers (16Smam1-To and 16Smam2; Table 1) containing Illumina sequencing adapters, 1 μM (5X) human blocking primer (16Smam-blkhum; Table1), 1 μM (5X) domestic pig *Sus scrofa* blocking primer (16Smam-blkpig; Table1), and 2 μL of direct PCR lysis buffer template solution from sample tubes for a final PCR reaction volume of 20 μL. PCR cycling conditions were: initial denaturation at 94 °C for 10 min, 40 cycles of 94 °C for 15 sec, 60 °C for 15 sec, and 68 °C for 20 sec, followed by a final elongation of 68 °C for 5 minutes. Each PCR run included a no-template control of sterile molecular grade water, a lysis buffer negative control, and a positive control containing DNA from bobcat *Lynx rufus*, a species that does not occur in South America. All negative and positive controls were indexed and sequenced.

**Table 1.**
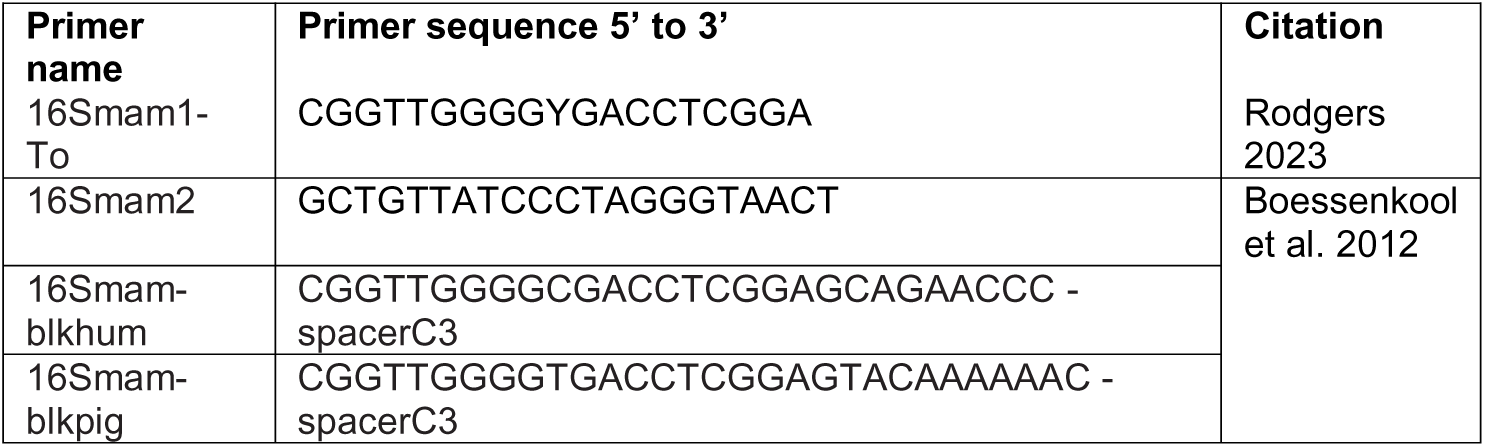
Primers used for metabarcoding of dung beetle and carrion fly-derived eDNA samples. The 1^st^ and 2^nd^ primers are forward and reverse primers respectively. The 3^rd^ and 4^th^ primers are blocking primers. These primers produce an amplicon of approximately 138 base pairs including primers.

PCR products from the initial PCR were diluted 50:1 in sterile H_2_O and then used as template for the indexing PCR. The indexing PCR was used to add unique dual indexes to each sample for multiplexing. Unique Dual Indexing (UDI) primers (Integrative DNA Technologies) contained the complimentary illumina flow-cell adapter sequence and an 8 bp unique dual index on each end (forward and reverse) for unique sample identification from multiplexed samples. Indexing PCRs reactions included 10 μl of Amplitaq Gold DNA polymerase (Thermo Fisher Scientific), 0.5 uM of forward and reverse UDI indexing primers, and 2 μl of diluted PCR product from the first PCR in a total reaction volume of 20 μl. PCR cycling conditions for the indexing PCR were: initial denaturation at 95 °C for 10 min, 15 cycles of 95 °C for 15 sec, 50 °C for 30 sec, and 72 °C for 30 sec, followed by a final elongation of 72 °C for 7 minutes. Following the indexing PCR, PCR product was cleaned up and normalized using a SequalPrep™ normalization plate (Thermo Fisher Scientific) following the manufacturer’s protocol. The resulting product was then pooled and provided to the Center for Integrative Biosystems sequencing core at Utah State University for sequencing on Illumina MiSeq.

PCR and sequencing were carried out on all fly samples. In addition, we tested two pooling methods for combining individual flies into pooled samples for sequencing (Fig 3). Because sequencing of individual flies may be prohibitively expensive for large datasets, we compared results from pooled samples with results from individual flies to see if pooling of samples led to loss of species detected. For the first method, we pooled 5 μL of lysis buffer from each of the 10 flies collected at each sampling site. Additionally, we pooled 5 μL of lysis buffer from each of the 40 flies collected from the same trail segment. The initial PCR was conducted using 4 μL of pools as template, after which the library preparation protocol was identical to that from individual flies. For the second pooling method, we pooled PCR product from the initial PCR of individual flies prior to indexing. For this method, we pooled 5 μL of PCR product from each of the 10 flies collected at each sampling site. Additionally, we made pools of PCR products from each of the 40 flies collected from the same trail, and from each of the 80 flies collected on each day of sampling. We then diluted this pooled PCR product 10:1 in sterile H_2_O and added 2 μL of pooled product to indexing PCR as above.

**Figure 3.**
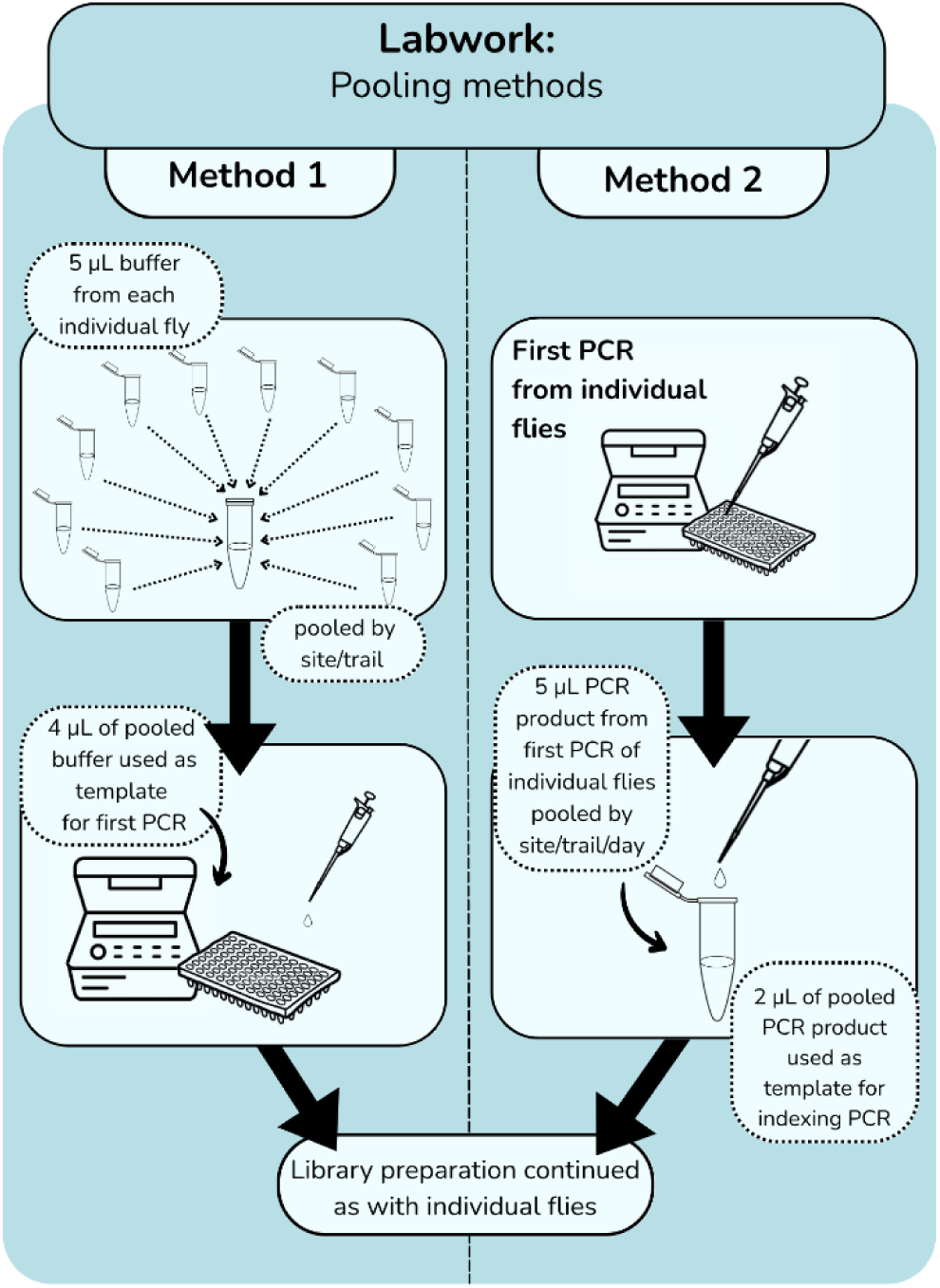
Two methods tested for pooling of samples to economize metabarcoding of iDNA from carrion flies for the detection of rainforest mammals from Los Amigos Biological Station, Madre de Dios, Peru.

Sequencing was carried out on three separate libraries. The first library contained 80 individual fly samples, plus 105 samples from a separate project. The second library contained the remaining 160 individual fly samples, plus samples from pooling method one. The third library contained only samples from pooling method two. All three libraries also contained positive and negative controls. Each library was sequenced on Illumina MiSeq with a 300 cycle v2 Micro reagent kit at the Center for Integrative Biosystems sequencing core at Utah State University.

Bioinformatic processing of MiSeq sequence data proceeded as follows. After demultiplexing, primer sequences were removed with CUTADAPT v. 1.18 (Martin, 2011). Next, data were filtered, denoised, paired-ends were merged, chimeras were removed with DADA2 (Callahan et al., 2016), and Amplicon Sequence Variants (ASV) were merged into Operational Taxonomic Units (OTUs) at 97% similarity with VSEARCH (Rognes et al., 2016), all within the QIIME2 environment (Bolyen et al., 2019) version 2023.5. For taxonomic assignment of OTUs, we compared two methods: BLAST (Altschul et al., 1990) and a Naïve Bayes classifier (Pedregosa et al., 2011) implemented within the QIIME2 environment. For BLAST, we retrieved the top 5 hits for each OTU, and taxonomy was assigned subjectively based on percent similarity of top hits (Table 2) with taxa known from the study area based on local expertise and a published species checklist from a nearby area (Payne et al., 2024). For taxonomic assignment with the Naïve Bayes classifier, we used the lrRNA (16S) reference database containing unique haplotypes from MIDORI 2 version GB265 (Leray et al., 2022). We first extracted our specific amplicon of interest from the database using our primer sequences, and then we trained a Naïve Bayes classifier using scikit-learn to assign taxonomy within QIIME2.

**Table 2.** Species detected from DNA metabarcoding of carrion flies from Los Amigos Biological Station, Madre de Dios, Peru, based on two taxonomic assignment methods, naïve bayes, and BLAST.

| Class | Order | Family | naïve bayes species | naïve bayes confidence | BLAST top hit species | BLAST match % | Presumed species | Species common name |
| --- | --- | --- | --- | --- | --- | --- | --- | --- |
| Mammalia | Artiodactyla | Cervidae | *Odocoileus sp.* | 0.857 | *Mazama americana* | 100.0 | *Mazama americana* | red brocket deer |
| Mammalia | Artiodactyla | Cervidae | *Passalites nemorivagus* | 0.937 | *Passalites nemorivagus* | 98.9 | *Passalites nemorivagus* | Amazonian brown brocket |
| Mammalia | Artiodactyla | Tayassuidae | *Catagonus wagneri* | 0.988 | *Catagonus wagneri* | 93.5 | *Tayassu pecari* | white-lipped peccary |
| Mammalia | Artiodactyla | Tayassuidae | *Dicotyles tajacu* | 1.000 | *Dicotyles tajacu* | 98.9 | *Dicotyles tajacu* | collared peccary |
| Mammalia | Carnivora | Felidae | *Puma concolor* | 1.000 | *Puma concolor* | 100.0 | *Puma concolor* | Puma |
| Mammalia | Carnivora | Mustelidae | *Unknown* | 1.000 | *Gulo gulo* | 96.7 | *Eira barbara* | Tyra |
| Mammalia | Carnivora | Procyonidae | *Bassaricyon alleni* | 0.984 | *Bassaricyon alleni* | 96.3 | *Bassaricyon alleni* | eastern lowland olingo |
| Mammalia | Carnivora | Procyonidae | *Nasua nasua* | 1.000 | *Nasua nasua* | 97.4 | *Nasua nasua* | South American coati |
| Mammalia | Carnivora | Procyonidae | *Potos flavus* | 1.000 | *Potos flavus* | 96.8 | *Potos flavus* | kinkajou |
| Mammalia | Chiroptera | Molossidae | *Molossus molossus* | 0.923 | *Molossus molossus* | 98.9 | *Molossus molossus* | velvety free-tailed bat |
| Mammalia | Chiroptera | Phyllostomidae | *Artibeus planirostris* | 0.988 | *Artibeus planirostris* | 100.0 | *Artibeus planirostris* | flat-faced fruit-eating bat |
| Mammalia | Chiroptera | Phyllostomidae | *Uroderma bilobatum* | 1.000 | *Uroderma bilobatum* | 96.7 | *Uroderma bilobatum* | tent-making bat |
| Mammalia | Cingulata | Dasypodidae | *Dasypus novemcinctus* | 0.788 | *Dasypus sabanicola* | 100.0 | *Dasypus sp.* | naked-tailed armadillo |
| Mammalia | Didelphimorphia | Didelphidae | *Didelphis marsupialis* | 0.919 | *Didelphis marsupialis* | 100.0 | *Didelphis marsupialis* | common opossum |
| Mammalia | Perissodactyla | Tapiridae | *Tapirus terrestris* | 0.971 | *Tapirus terrestris* | 100.0 | *Tapirus terrestris* | Lowland tapir |
| Mammalia | Primates | Aotidae | *Aotus sp.* | 1.000 | *Aotus nigriceps* | 100.0 | *Aotus nigriceps* | black-headed night monkey |
| Mammalia | Primates | Atelidae | *Alouatta seniculus* | 1.000 | *Alouatta seniculus* | 100.0 | *Alouatta seniculus* | Red howler |
| Mammalia | Primates | Atelidae | *Ateles chamek* | 0.977 | *Ateles chamek* | 100.0 | *Ateles chamek* | black spider monkey |
| Mammalia | Primates | Cebidae | *Cebuella sp.* | 0.974 | *Cebuella pygmaea* | 93.4 | *Cebuella niveiventris* | Eastern pygmy marmoset |
| Mammalia | Primates | Cebidae | *Cebus unicolor* | 0.999 | *Cebus unicolor* | 100.0 | *Cebus cuscinus* | Shock-headed capuchin |
| Mammalia | Primates | Cebidae | *Leontocebus fuscicollis* | 1.000 | *Leontocebus fuscicollis weddelli* | 100.0 | *Leontocebus weddelli* | Weddell's saddle-back tamarin |
| Mammalia | Primates | Cebidae | *Saimiri sp.* | 1.000 | *Saimiri boliviensis* | 98.9 | *Saimiri boliviensis* | Bolivian squirrel monkey |
| Mammalia | Primates | Cebidae | *Sapajus* | 1.000 | *Sapajus apella* | 100.0 | *Sapajus apella* | tufted capuchin |
| Mammalia | Rodentia | Cricetidae | *Oligoryzomys flavescens* | 0.930 | *Oligoryzomys flavescens* | 86.4 | *Oligoryzomys sp.* | pygmy rice rat |
| Mammalia | Rodentia | Cuniculidae | *Cuniculus paca* | 1.000 | *Cuniculus paca* | 100.0 | *Cuniculus paca* | Lowland paca |
| Mammalia | Rodentia | Echimyidae | *Mesomys hispidus* | 0.985 | *Mesomys hispidus* | 95.7 | *Mesomys hispidus* | spiny tree rat |
| Mammalia | Rodentia | Erethizontidae | *Coendou insidiosus* | 0.999 | *Coendou insidiosus* | 94.6 | *Coendou sp.* | prehensile-tailed porcupine |
| Amphibia | Anura | Hylidae | *Dendropsophus koechlini* | 1.000 | *Dendropsophus koechlini* | 100.0 | *Dendropsophus koechlini* | Koechlin's treefrog |
| Amphibia | Anura | Hylidae | *Scinax ictericus* | 1.000 | *Scinax ictericus* | 100.0 | *Scinax ictericus* | Tree frog species |
| Amphibia | Anura | Leptodactylidae | *Leptodactylus stenodema* | 0.999 | *Leptodactylus stenodema* | 91.8 | *Leptodactylus stenodema* | San Jose white-lipped frog |
| Amphibia | Anura | Microhylidae | *Ctenophryne geayi* | 1.000 | *Ctenophryne geayi* | 100.0 | *Ctenophryne geayi* | brown egg frog |

Because pooling method two had approximately 9 times greater sequencing read-depth than pooling method one (see results), we also conducted a rarefication of reads in pooling method two samples to see if rarefaction changed results. Rarefaction was carried out in QIIME2, with a reduced sampling depth of 10,245 reads per sample to match read-depth of pooling method one samples.

For OTUs that could not be assigned with high confidence at the species level with either of our classification methods, in some cases we subjectively assigned a “presumed species” based on local expertise of the known mammal community, and a species checklist from a nearby area (Payne et al., 2024). This was done when an OTU was assigned only to genus with confidence, but when only one species in the given genus is known from the study area. In two cases, we also subjectively assigned “presumed species” to OTUs only assigned at the family level, but that we believe belong to taxa from our study area that lack reference sequences (Table 2).

**Figure 4.**
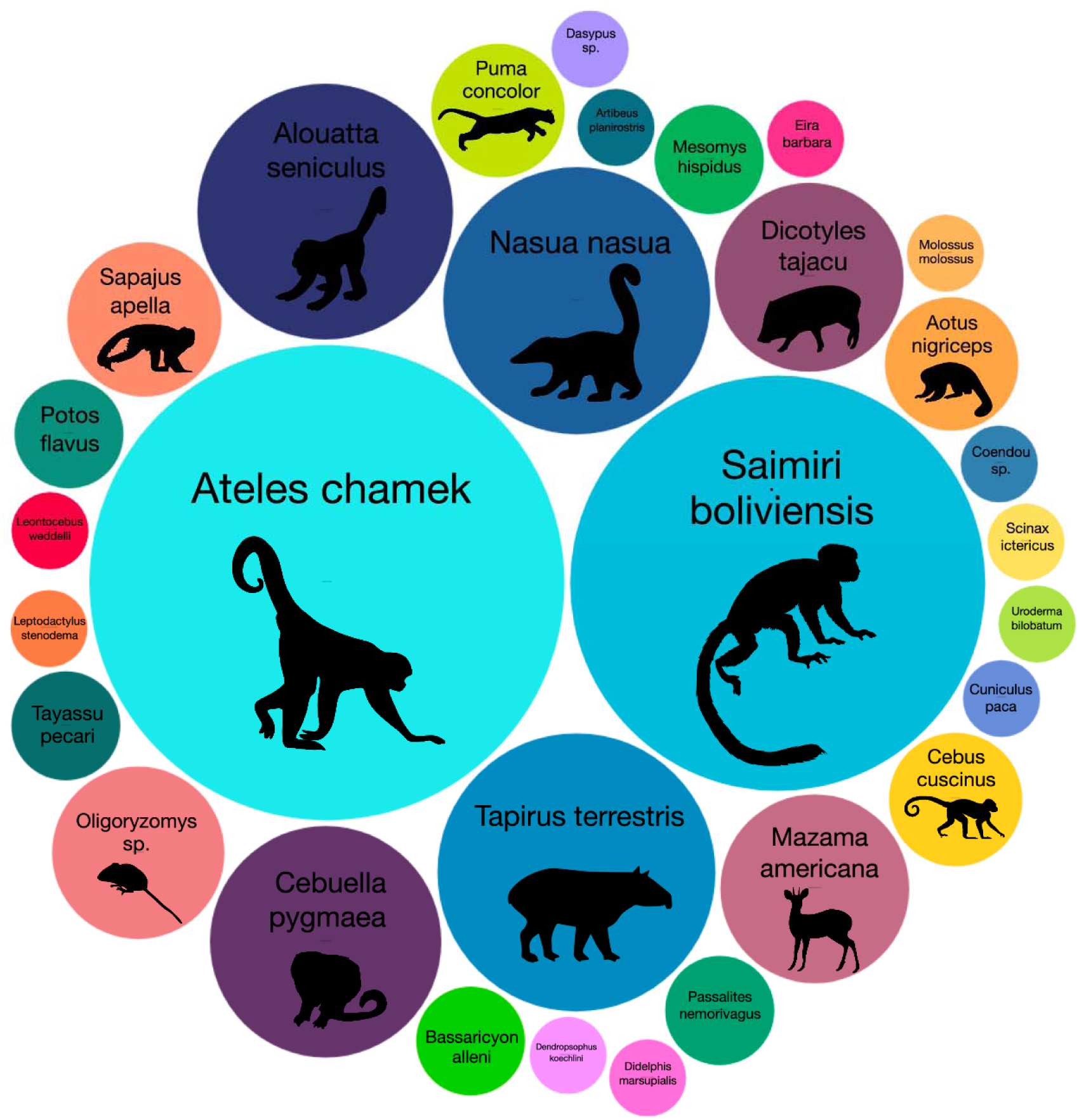
Visual representation of mammals detected by metabarcoding of carrion flies from Los Amigos Biological Station, Madre de Dios, Peru. Size of circles represents the percentage of flies containing vertebrate DNA for each species.

## Results

### Carrion fly collection

Two hundred and forty carrion flies were collected from 24 locations along six trails. Because of the difficulty of identifying carrion flies to species using morphology, flies were in most cases only identified to family. The most common families collected were Calliphoridae, Sarcophagidae, Mesembrinellidae, and Tachinidae representing 53%, 35%, 6%, and 3% of total flies collected respectively. A few flies were also collected from the families Syrphidae (n=3) and Muscidea (n=3).

### Carrion fly laboratory analysis

MiSeq sequencing of carrion fly samples from the first two libraries (individual flies and pool samples from pooling method one) resulted in 3,965,439 reads, of which 2,547,579 reads were retained after quality filtering, denoising, and chimera removal, with a median of 9,956 reads per individual fly sample, and a median of 10,245 reads per pool sample. MiSeq sequencing of carrion fly pool samples from the third library (pooling method two) resulted in 3,280,546 reads, of which 3,086,005 reads were retained after quality filtering, denoising, and chimera removal, with a median of 92,385 reads per pool sample. These reads were clustered into 78 unique OTUs. After removal of contaminant OTUs identified by BLAST as belonging to human or domestic species and the *Lynx rufus* positive control, and OTUs that could not be identified to the family level, 40 unique OTUs assigned to vertebrate species remained.

These 40 OTUs were assigned to 31 unique taxa including eight primates, five carnivores (one felid, three procyonids, and one mustelid), four rodents, four artiodactyls, one perissodactyl, three chiropterans, one armadillo, one marsupial, and four anurans (Table 2). Of these 31 unique taxa, 26 were assigned to a species known from the study area. Of the remaining 5 taxa, 3 were assigned only at the genus level, and 2 at the family level. We subjectively assigned a “presumed species” to two OTUs that our classifiers could only assign objectively to the family level. In both cases, a reliable reference sequence for 16S for the species in question was not available in GenBank. However, in both cases, all other species from the given family known from the study area do possess a reliable reference sequence, leading us to believe that these OTUs belong to the “presumed species”. In one case, an OTU was identified by both classifiers as Chacoan peccary *Catagonus wagneri* (Table 2). The Chacoan peccary does not occur in Peru, and thus, it is not possible that the sequence originated from this species. However, the white lipped peccary *Tayassu pecari* is present at Los Amigos, but a reliable 16S reference sequence for this species is not available in GenBank. GenBank does possess a partial mitochondria genome for *T. pecari* (Accession: KY987556), but this genome is low quality, with many ambiguous bases in the region corresponding to our amplicon. *C. wagneri* is the most closely related extant species to *T. pecari* and is more closely related to *T. pecari* than to collared peccary *Pecari tajacu* (Parisi Dutra et al., 2017), the other peccary species present at Los Amigos. Thus, we presume this OTU belongs to white lipped peccary. In another case, one OTU was assigned by the Naïve Bayes classifier only to the family Mustelidae, while BLAST returned a top hit at 96.7% to *Gulo gulo* or wolverine, with all 5 top hits assigned to species within the family Mustelidae, but none to species known from Los Amigos. Three of the four mustelid species known from Los Amigos have a 16S reference sequence available apart from the tayra *Eira barbara*. Thus, we presume this OTU belongs to *E. barbara.* In total, we detected and taxonomically assigned vertebrate DNA from 106/240 (44.2%) individual flies (excluding human or domestic species). Among these, 63 flies yielded DNA from a single species, 31 flies yielded DNA from two species, ten flies yielded DNA from three species, one fly yielded DNA from four species, and one fly yielded DNA from five species. The mean number of flies yielding DNA per site was 4.0 (range 1-6). The mean number of species detected per site was 4.96 (range 1-8), with at least one vertebrate species detected from all 24 sites.

Of the 106 fly samples that contained vertebrate DNA, 67 (63%) were from the family Calliphoridea, 31 (29%) were from the family Sarcopagidea, three (2.8%) were from the family Tachinidae, three (2.8%) were from the family Mesembrinellidae, and two (1.8%) were from the family Muscidae. This corresponds to 53% (67/126) of flies from the family Calliphoridea, 37% (31/84) of flies from the family Sarcopagidea, 37% (3/8) of the flies from the family Tachinidae, 67% (2/3) of the flies from the family Muscidae, and 20% (3/15) of the flies from the family Mesembrinellidae containing vertebrate DNA. No vertebrate DNA was detected from the three flies from the family Syrphidae.

Primates were the most commonly detected mammalian order. Primate DNA was detected from 71/240 (29.6%) of all flies collected, with Black spider monkey *Ateles chamek* detected from 40/240 flies (16.67%) and Bolivian squirrel monkey *Saimiri boliviensis* detected from 31/240 flies (12.92%; Table 3). Lowland tapir *Tapirus terrestris* was the third most commonly detected species and was detected from 14/240 flies (5.83%; Table 3).

**Table 3.**
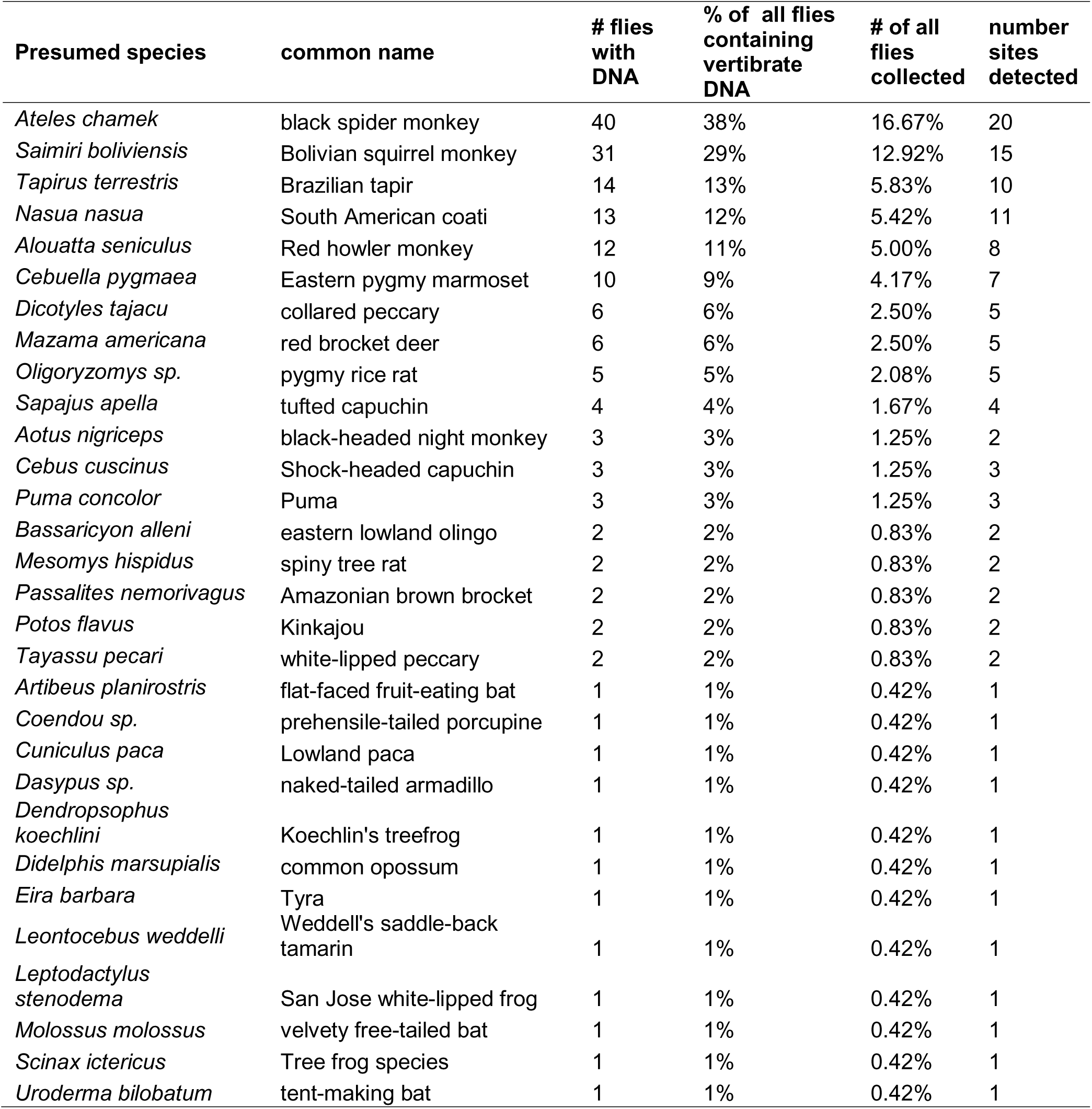
Summary of the number and percentage of flies collected containing DNA from each presumed species from metabarcoding of DNA from carrion flies collected from Los Amigos Biological Station, Madre de Dios, Peru.

### Fly pooling methods

Pooling method one (pooling of buffer prior to initial PCR) resulted in detection of a mean of 2.25 (range 0-5) species per site (n=10 flies per pool) compared to a mean of 4.96 (range 1-8) species per site from individual flies. Pooling method one detected a mean of 4.0 (range 1-7) vertebrate species per trail segment (n=40 flies per pool) compared to a mean of 12 species per trail segment from individual flies (Table S1). Pooling method one detected the same species detected from individual flies at just 1/24 sites (4.2%), and at 0/6 trail segments.

Pooling method two, (pooling of PCR product after initial PCR from individual flies and prior to indexing) resulted in detection of a mean of 5.04 (range 1-8) species per site (n=10 flies per pool), just slightly more than the mean number of species detected from individual flies (4.96 species per site). At two sites, individual flies detected one more species than the PCR product pool. At the 22 remaining sites, the PCR pool detected the exact same species detected by individual flies. At three sites, the PCR product pool detected one additional species not detected by individual flies, and at one site, the pool detected two additional species. Pooling Method two detected a mean of 12.33 species per trial segment (n=40 flies per pool), slightly higher than the mean number of species detected from individual flies (12 species). This method detected a mean of 17 species per day (n=80 flies per pool), slightly higher than the mean number of species detected from individual flies (16.67 species; Table S1).

After rarifying read-depth from pooling method two samples to match depth of pooling method one samples, results remained nearly identical to the results from the non-rarified dataset. Two exceptions were noted: at one site, *Cebuella niveiventris* was detected from the non-rarified dataset, but not from the rarified dataset, and at another site, *Dendropsophus koechlini* was detected from the non-rarified dataset, but not from the rarified dataset. Otherwise, the results were identical between the rarified and non-rarified datasets.

## Discussion

Our results clearly demonstrate that carrion fly-derived iDNA is an efficient method for surveying mammal species from tropical forests. Twenty-seven unique mammalian taxa from 9 orders and 18 families plus four anuran taxa from three families were detected from just 240 carrion flies collected over the course of just 3 days. This methodology was highly effective at detecting arboreal species which are often difficult to detect with camera traps, including eight primates, among them the near-threatened shock-headed capuchin *Cebus cuscinus,* the vulnerable eastern pygmy marmoset *Cebuella niveiventris,* and the endangered black spider monkey *Ateles chamek* (IUCN, 2025).

A key methodological advancement of this project was the use of an extraction-free, direct PCR approach to target DNA left by carrion flies in collection tubes. While we are not the first to use this approach for carrion flies (see Srivathsan et al., 2023), we found that this innovative approach offers significant advantages; it not only streamlines fieldwork but also reduces laboratory costs by eliminating the labor-intensive DNA extraction step. Previous approaches for carrion fly metabarcoding required grinding whole flies into a slurry for DNA extraction (Calvignac-Spencer, Merkel, et al., 2013; T. W. Rodgers et al., 2017) or dissecting fly digestive tracts (Lee et al., 2016), processes that demand substantial effort. In contrast, the extraction-free method uses a simple buffer added to the empty tubes that previously contained carrion flies as DNA template in PCR, bypassing the need for grinding or dissecting flies followed by DNA extraction. This method saves significant time, effort, and cost. Moreover, flies could be released alive after depositing DNA into the tubes (although ours were preserved for taxonomic identification), making this method potentially even less invasive, with minimal impact on fly populations. This extraction-free, direct PCR approach significantly improves the speed and efficiency of iDNA techniques for surveying and monitoring vertebrate biodiversity.

Another goal of this study was to improve the scalability of using carrion flies for large-scale surveys of vertebrate diversity. Although the current study used just 240 carrion flies collected over three days, future studies may utilize thousands of flies collected over much larger areas and timescales. To this end, we tested a method for collecting carrion flies by netting them directly off a bag of bait rather than using bottle-traps as has been done in some past studies (e.g. Rodgers et al., 2017). This method proved to be effective, as we were able to collect 10 flies from each site in approximately 15-20 minutes. This method is advantageous compared to trapping with bottle-traps for two reasons. First, it only requires visiting each site one time to collect flies, whereas trapping requires moving traps from location to location, and then visiting the traps the following day to collect flies. Second, it requires very little equipment, just a good insect net, and a plastic bag full of bait. This is highly preferable for large-scale studies sampling from many locations, as each site can be visited just once, for a short period of time, without the need to return to collect traps.

Two different methods were evaluated for metabarcoding of pooled samples containing multiple carrion flies. This provides important insight because for larger-scale studies with potentially thousands of individual flies it would be prohibitively expensive to sequence each fly individually. The ability to pool flies collected from the same site prior to sequencing is advantageous to decrease sequencing cost while still maintaining the ability to differentiate between sites. For instance, Rodgers et al. (2017) pooled 16 ground-up flies into each DNA extraction but did not sequence individual flies for comparison. Pooling of multiple flies, however, may lead to missed species detections due to sample dilution. To this end, we tested two methods for pooling fly samples that were collected at the same site. Pooling method one involved pooling buffers from flies collected from the same site prior to PCR, and using the pooled buffer as template for PCR. This method resulted in the loss of many species compared to sequencing of individual flies, likely due to sample dilution such that template DNA from all species was not present in the aliquot added to PCR, and thus not amplified.

Conversely, pooling method two, pooling of PCR products after conducting initial PCR from individual fly samples, but prior to sample indexing, performed exceptionally well. Minor discrepancies were noted, but overall, the results from this second method were nearly identical to results obtained from individual flies. This method is more time-consuming and expensive than pooling method one, as an initial PCR must be conducted from each fly sample. However, this method would still allow many more fly samples to be multiplexed and sequenced in a single Next-Generation Sequencing run, making it a feasible method for large-scale studies with large numbers of samples. Initial sequencing depth was approximately nine times higher for pooling method two versus method one. As this difference in read-depth could have accounted for some of the observed discrepancy between the two methods tested, we also rarified read-depth from pooling method two samples to match depth from pooling method one samples, to ensure that differences in species detection between the two methods were not due to differences in read-depth. Results from the rarified dataset were nearly identical to results from the non-rarified dataset, indicating that read-depth clearly did not account for the differences in species detection seen between the two methods. Thus, method two is clearly preferable, as it performed nearly identically to sequencing of individual flies, without loss of species, while method one resulted in the loss of many species, even though it is possible results from method one may have improved with greater sequencing depth.

We found that, in a few cases, species were detected in sample pools that were not detected from individual flies. Because PCR from individual flies were run with just 2 μL (5%) of the total buffer collected from each tube as template, it is likely that in some cases, DNA from a species was present in the tube, but was not present in the template that was added to PCR. For example, the brown egg frog *Ctenophryne geayi* was detected from pooling method one but was not detected by individual flies or by pooling method two (Table S1). Only 49 reads of this species were detected from a single pool sample indicating that the DNA of this species was likely at low concentration. Running a greater number of PCR replicates per sample would decrease the chance of missing species with very little DNA present, however this would substantially increase cost. Another potential approach would be to pool buffer and then concentrate the buffer pool such that the entire pool could be included as template in a single PCR. This approach could potentially be more cost effective and should be investigated in future research.

We tested two different methods for taxonomic assignment of OTUs derived from carrion fly metabarcoding, BLAST (Altschul et al., 1990) and a Naive Bayes classifier using scikit-learn (Pedregosa et al., 2011) implemented within the QIIME2 environment. Both methods provided consistent assignments for nearly all OTUs with a few exceptions (Table 2). In one case the Naïve Bayes classifier classified an OTU at a relatively low confidence of 0.86 to the genus *Odocoileus*, a genus of cervid that is not known from the study area. BLAST returned a hit for *Mazama americana*, a cervid species that is known from our study area, with a 100% match to the reference sequence. Thus, we presumed this OTU originated from *M. americana.* Interestingly, however, BLAST also returned hits that were a 100% match to *Odocoileus virginianus, Odocoileus hemionus, Mazama nana, Mazama rufa,* and *Mazama bororo,* none of which occur in our study area. Thus, while we still conclude this OTU likely originated from *M. americana*, the 16S marker alone is insufficient in differentiating between these species. In another case, both methods identified one OTU as *Passalites nemorivagus*, the Amazonian brown brocket, a species which is known from our study area. This OTU was a 98.9% match to a reference sequence from *P. nemorivagus* but was also a 98.9% match to reference sequences from *Ozotoceros bezoarticus*, a South American cervid species not known from our study area, and *Rangifer tarandus* or Reindeer. Thus, it is likely that the 16S amplicon used in this study may have relatively poor taxonomic power for distinguishing cervid species. These cases emphasize the fact that, in many instances, prior knowledge of the local wildlife community can aid in the correct assignment of taxa at the species level. Additionally, using multiple taxonomic classifiers can assist the assignment of sequences to the correct species. In the case of the OTU ultimately assigned to *M. americana,* use of only the Naïve Bayes classifier would have led to an incorrect assignment. We also potentially detected two mammal species from our study area that lack reference sequences for the 16S amplicon used in our study: *Tayassu pecari*, and *Eira barbara.* As these species have wide Neotropical distributions and are of conservation concern in some areas, generating 16S reference sequences for these species should be a future priority so that they can be accurately identified in future eDNA/iDNA studies.

In two instances, sequences were assigned to primate species that are not known from our study area. One OTU was assigned with high probability by both classifiers as the capuchin monkey species *Cebus unicolor. C. unicolor* is not known from our study area, however a closely related species, *C. cuscinus* is. *C. cuscinus* and *C. unicolor* were historically considered subspecies of *C. albifrons,* however they have since been split into separate species (Boubli et al., 2012). Although this split was based in part on mitochondrial genetics, it appears likely that *C. cuscinus* and *C. unicolor* are genetically identical at our short 16S mitochondrial locus, as the sequence we found was a 100% match to a *C. unicolor* sequence. No *C. cuscinus* reference sequences are available, however we presume that the sequence we detected is from *C. cuscinus,* as this is the only *Cebus* species known from our study area. Both classifiers also assigned an OTU to the genus *Cebuella* or pygmy marmoset. Pygmy marmosets are not known to be present at Los Amigos, however, the eastern pygmy marmoset *Cebuella niveiventris* is known from the region. Both classifiers identified this sequence with somewhat lower confidence, the naïve bayes classifier only identified the sequence to genus level at 0.974 confidence, and BLAST had a top hit match percentage of just 93.4% with the sister species *Cebuella pygmaea.* Thus, it is possible that this classification is artifactual. However, it would seem unlikely that a sequence would be artificially assigned to a species known from the region just by chance. This sequence was also found in 10 individual flies, making it the 6^th^ most commonly detected species in our study, which is peculiar for a species that has never been observed in our study area. One possibility is that a dead marmoset floated downriver from a population upriver and was being actively consumed by flies when our sampling occurred, as this would explain why the sequence was detected from a relatively large number of flies. Because this detection is unexpected, and confidence in the assignment were lower than for most other species, it should be treated with some uncertainty.

In conclusion, this study demonstrates the effectiveness of using carrion fly-derived iDNA for surveying mammalian assemblages in tropical rainforest. We detected 28 unique mammal species plus four anuran species from just 240 carrion flies. A key methodological advancement was the extraction-free, direct PCR approach we used, which simplifies both field and lab work, significantly reducing time and costs while maintaining robust species detection. This method offers a non-invasive and efficient approach for rainforest mammal monitoring and appears to be particularly valuable for the detection of arboreal primates. Additionally, we investigated methods for scaling-up carrion fly metabarcoding surveys, including a more efficient collection strategy, as well as pooling methods for cost-effective sequencing. Pooling PCR products after initial amplification from individual flies, but prior to indexing, proved highly effective, nearly perfectly matching results from sequencing of individual flies without sacrificing species detection accuracy, making it a viable method for future large-scale iDNA surveys. Our findings will contribute to the efficiency and practicability of using iDNA metabarcoding for future biodiversity monitoring and conservation of mammals in tropical forests.

## Acknowledgments

We thank the staff at Los Amigos Biological Station for making this work possible. We thank Sam Pottie for contributing expertise on Amazonian primate species. Funding for this work was provided by the Gordon and Betty Moore Foundation. This work was conducted on the ancestral lands of Indigenous peoples of the Madre de Dios region, including the Yine, Matsigenka, Harakbut, and others. We recognize their enduring cultural, spiritual, and ecological stewardship of these forests and the tremendous biodiversity they contain, and we honor their sovereignty and knowledge.

## Notes

### Competing Interest Statement

The authors have declared no competing interest.

