## Supplementary material for "An efficient iDNA method for surveying rainforest mammals with carrion flies": Table S1

**Supplementary Information**

**Table S1**. Vertebrate species detected from different pooling methods from iDNA metabarcoding of carrion flies from Los Amigos Biological Station, Peru.

| **Site** | **Species detected individual flies** | **# Species** | **Species detected Pool 1** |  | **Species detected Pool 2** | **# Species** |
| --- | --- | --- | --- | --- | --- | --- |
| Site 1 | *Ateles chamek* | 6 | *Nasua nasua* | 1 | *Ateles chamek* | 6 |
|  | *Bassaricyon alleni* |  |  |  | *Bassaricyon alleni* |  |
|  | *Cuniculus paca* |  |  |  | *Cuniculus paca* |  |
|  | *Nasua nasua* |  |  |  | *Nasua nasua* |  |
|  | *Oligoryzomys sp.* |  |  |  | *Oligoryzomys sp.* |  |
|  | *Saimiri boliviensis* |  |  |  | *Saimiri boliviensis* |  |
| Site 2 | *Oligoryzomys sp.* | 1 |  | 0 | *Oligoryzomys sp.* | 1 |
| Site 3 | *Ateles chamek* | 3 | *Ateles chamek* | 1 | *Ateles chamek* | 4 |
|  | *Dasypus sp.* |  |  |  | *Dasypus sp.* |  |
|  | *Passalites nemorivagus* |  |  |  | *Passalites nemorivagus* |  |
|  |  |  |  |  | *Sapajus apella* |  |
| Site 4 | *Ateles chamek* | 3 |  | 0 | *Ateles chamek* | 3 |
|  | *Dicotyles tajacu* |  |  |  | *Dicotyles tajacu* |  |
|  | *Tapirus terrestris* |  |  |  | *Tapirus terrestris* |  |
| Site 5 | *Ateles chamek* | 5 | *Ateles chamek* | 5 | *Ateles chamek* | 5 |
|  | *Cebuella niveiventris* |  | *Cebuella niveiventris* |  | *Cebuella niveiventris* |  |
|  | *Leontocebus weddelli* |  | *Leontocebus weddelli* |  | *Leontocebus weddelli* |  |
|  | *Nasua nasua* |  | *Nasua nasua* |  | *Nasua nasua* |  |
|  | *Saimiri boliviensis* |  | *Saimiri boliviensis* |  | *Saimiri boliviensis* |  |
| Site 6 | *Ateles chamek* | 5 | *Ateles chamek* | 4 | *Ateles chamek* | 7 |
|  | *Nasua nasua* |  | *Cebus cuscinus* |  | *Cebus cuscinus* |  |
|  | *Puma concolor* |  | *Saimiri boliviensis* |  | *Nasua nasua* |  |
|  | *Saimiri boliviensis* |  | *Sapajus apella* |  | *Puma concolor* |  |
|  | *Tapirus terrestris* |  |  |  | *Saimiri boliviensis* |  |
|  |  |  |  |  | *Sapajus apella* |  |
|  |  |  |  |  | *Tapirus terrestris* |  |
| Site 7 | *Ateles chamek* | 5 | *Dicotyles tajacu* | 3 | *Ateles chamek* | 5 |
|  | *Dicotyles tajacu* |  | *Saimiri boliviensis* |  | *Dicotyles tajacu* |  |
|  | *Saimiri boliviensis* |  | *Tapirus terrestris* |  | *Saimiri boliviensis* |  |
|  | *Tapirus terrestris* |  |  |  | *Tapirus terrestris* |  |
|  | *Tayassu pecari* |  |  |  | *Tayassu pecari* |  |
| Site 8 | *Nasua nasua* | 3 | *Passalites nemorivagus* | 1 | *Nasua nasua* | 3 |
|  | *Passalites nemorivagus* |  |  |  | *Passalites nemorivagus* |  |
|  | *Tapirus terrestris* |  |  |  | *Tapirus terrestris* |  |
| Site 9 | *Alouatta seniculus* | 4 | *Alouatta seniculus* | 3 | *Alouatta seniculus* | 4 |
|  | *Ateles chamek* |  | *Ateles chamek* |  | *Ateles chamek* |  |
|  | *Nasua nasua* |  | *Tapirus terrestris* |  | *Nasua nasua* |  |
|  | *Tapirus terrestris* |  |  |  | *Tapirus terrestris* |  |
| Site 10 | *Ateles chamek* | 6 | *Ateles chamek* | 4 | *Ateles chamek* | 7 |
|  | *Bassaricyon alleni* |  | *Cebuella niveiventris* |  | *Bassaricyon alleni* |  |
|  | *Cebuella niveiventris* |  | *Nasua nasua* |  | *Cebuella niveiventris* |  |
|  | *Nasua nasua* |  | *Potos flavus* |  | *Nasua nasua* |  |
|  | *Potos flavus* |  |  |  | *Potos flavus* |  |
|  | *Puma concolor* |  |  |  | *Puma concolor* |  |
|  |  |  |  |  | *Tapirus terrestris* |  |
| Site 11 | *Artibeus planirostris* | 7 | *Artibeus planirostris* | 2 | *Artibeus planirostris* | 6 |
|  | *Ateles chamek* |  | *Ateles chamek* |  | *Ateles chamek* |  |
|  | *Dicotyles tajacu* |  |  |  | *Mesomys hispidus* |  |
|  | *Mesomys hispidus* |  |  |  | *Saimiri boliviensis* |  |
|  | *Saimiri boliviensis* |  |  |  | *Tapirus terrestris* |  |
|  | *Tapirus terrestris* |  |  |  | *Tayassu pecari* |  |
|  | *Tayassu pecari* |  |  |  |  |  |
| Site 12 | *Alouatta seniculus* | 8 | *Alouatta seniculus* | 2 | *Alouatta seniculus* | 8 |
|  | *Aotus nigriceps* |  | *Potos flavus* |  | *Aotus nigriceps* |  |
|  | *Ateles chamek* |  |  |  | *Ateles chamek* |  |
|  | *Coendou sp.* |  |  |  | *Coendou sp.* |  |
|  | *Oligoryzomys sp.* |  |  |  | *Oligoryzomys sp.* |  |
|  | *Potos flavus* |  |  |  | *Potos flavus* |  |
|  | *Saimiri boliviensis* |  |  |  | *Saimiri boliviensis* |  |
|  | *Sapajus apella* |  |  |  | *Sapajus apella* |  |
| Site 13 | *Aotus nigriceps* | 5 | *Aotus nigriceps* | 2 | *Aotus nigriceps* | 4 |
|  | *Leptodactylus stenodema* |  | *Saimiri boliviensis* |  | *Mazama americana* |  |
|  | *Mazama americana* |  |  |  | *Saimiri boliviensis* |  |
|  | *Saimiri boliviensis* |  |  |  | *Tapirus terrestris* |  |
|  | *Tapirus terrestris* |  |  |  |  |  |
| Site 14 | *Ateles chamek* | 6 | *Ateles chamek* | 2 | *Ateles chamek* | 6 |
|  | *Cebuella niveiventris* |  | *Saimiri boliviensis* |  | *Cebuella niveiventris* |  |
|  | *Dicotyles tajacu* |  |  |  | *Dicotyles tajacu* |  |
|  | *Mazama americana* |  |  |  | *Mazama americana* |  |
|  | *Saimiri boliviensis* |  |  |  | *Saimiri boliviensis* |  |
|  | *Uroderma bilobatum* |  |  |  | *Uroderma bilobatum* |  |
| Site 15 | *Alouatta seniculus* | 6 | *Alouatta seniculus* | 4 | *Alouatta seniculus* | 6 |
|  | *Ateles chamek* |  | *Ateles chamek* |  | *Ateles chamek* |  |
|  | *Cebuella niveiventris* |  | *Passalites nemorivagus* |  | *Cebuella niveiventris* |  |
|  | *Mazama americana* |  | *Puma concolor* |  | *Mazama americana* |  |
|  | *Puma concolor* |  |  |  | *Puma concolor* |  |
|  | *Saimiri boliviensis* |  |  |  | *Saimiri boliviensis* |  |
| Site 16 | *Ateles chamek* | 3 | *Ateles chamek* | 1 | *Ateles chamek* | 3 |
|  | *Molossus molossus* |  |  |  | *Molossus molossus* |  |
|  | *Sapajus apella* |  |  |  | *Sapajus apella* |  |
| Site 17 | *Ateles chamek* | 3 |  | 0 | *Ateles chamek* | 3 |
|  | *Didelphis marsupialis* |  |  |  | *Didelphis marsupialis* |  |
|  | *Mesomys hispidus* |  |  |  | *Mesomys hispidus* |  |
| Site 18 | *Alouatta seniculus* | 6 | *Alouatta seniculus* | 4 | *Alouatta seniculus* | 6 |
|  | *Ateles chamek* |  | *Ateles chamek* |  | *Ateles chamek* |  |
|  | *Cebuella niveiventris* |  | *Nasua nasua* |  | *Cebuella niveiventris* |  |
|  | *Nasua nasua* |  | *Tapirus terrestris* |  | *Nasua nasua* |  |
|  | *Saimiri boliviensis* |  |  |  | *Saimiri boliviensis* |  |
|  | *Tapirus terrestris* |  |  |  | *Tapirus terrestris* |  |
| Site 19 | *Ateles chamek* | 4 | *Ateles chamek* | 1 | *Ateles chamek* | 4 |
|  | *Cebuella niveiventris* |  |  |  | *Cebuella niveiventris* |  |
|  | *Oligoryzomys sp.* |  |  |  | *Oligoryzomys sp.* |  |
|  | *Saimiri boliviensis* |  |  |  | *Saimiri boliviensis* |  |
| Site 20 | *Alouatta seniculus* | 5 | *Alouatta seniculus* | 4 | *Alouatta seniculus* | 5 |
|  | *Ateles chamek* |  | *Ateles chamek* |  | *Ateles chamek* |  |
|  | *Cebus cuscinus* |  | *Mazama americana* |  | *Cebus cuscinus* |  |
|  | *Oligoryzomys sp.* |  | *Saimiri boliviensis* |  | *Oligoryzomys sp.* |  |
|  | *Saimiri boliviensis* |  |  |  | *Saimiri boliviensis* |  |
| Site 21 | *Alouatta seniculus* | 6 | *Ateles chamek* | 1 | *Alouatta seniculus* | 6 |
|  | *Ateles chamek* |  |  |  | *Ateles chamek* |  |
|  | *Mazama americana* |  |  |  | *Mazama americana* |  |
|  | *Nasua nasua* |  |  |  | *Nasua nasua* |  |
|  | *Saimiri boliviensis* |  |  |  | *Saimiri boliviensis* |  |
|  | *Sapajus apella* |  |  |  | *Sapajus apella* |  |
| Site 22 | *Alouatta seniculus* | 7 | *Alouatta seniculus* | 6 | *Alouatta seniculus* | 8 |
|  | *Ateles chamek* |  | *Ateles chamek* |  | *Ateles chamek* |  |
|  | *Cebuella niveiventris* |  | *Cebuella niveiventris* |  | *Cebuella niveiventris* |  |
|  | *Eira barbara* |  | *Nasua nasua* |  | *Dendropsophus koechlini* |  |
|  | *Nasua nasua* |  | *Saimiri boliviensis* |  | *Eira barbara* |  |
|  | *Saimiri boliviensis* |  | *Tapirus terrestris* |  | *Nasua nasua* |  |
|  | *Tapirus terrestris* |  |  |  | *Saimiri boliviensis* |  |
|  |  |  |  |  | *Tapirus terrestris* |  |
| Site 23 | *Alouatta seniculus* | 8 | *Oligoryzomys sp.* | 2 | *Alouatta seniculus* | 8 |
|  | *Ateles chamek* |  | *Saimiri boliviensis* |  | *Ateles chamek* |  |
|  | *Cebus cuscinus* |  |  |  | *Cebus cuscinus* |  |
|  | *Mazama americana* |  |  |  | *Mazama americana* |  |
|  | *Nasua nasua* |  |  |  | *Nasua nasua* |  |
|  | *Saimiri boliviensis* |  |  |  | *Saimiri boliviensis* |  |
|  | *Sapajus apella* |  |  |  | *Sapajus apella* |  |
|  | *Scinax ictericus* |  |  |  | *Scinax ictericus* |  |
| Site 24 | *Cebus cuscinus* | 3 | *Nasua nasua* | 1 | *Cebus cuscinus* | 3 |
|  | *Dicotyles tajacu* |  |  |  | *Dicotyles tajacu* |  |
|  | *Nasua nasua* |  |  |  | *Nasua nasua* |  |
| **Mean # species per site** |  | **4.96** |  | **2.25** |  | **5.04** |
| Trail seg. M1 | Ateles chamek | 10 | *Bassaricyon alleni* | 2 | *Ateles chamek* | 11 |
|  | Bassaricyon alleni |  | *Nasua nasua* |  | *Bassaricyon alleni* |  |
|  | Cuniculus paca |  |  |  | *Cuniculus paca* |  |
|  | Dasypus sp. |  |  |  | *Dasypus sp.* |  |
|  | Dicotyles tajacu |  |  |  | *Dicotyles tajacu* |  |
|  | Nasua nasua |  |  |  | *Nasua nasua* |  |
|  | Oligoryzomys sp. |  |  |  | *Oligoryzomys sp.* |  |
|  | Passalites nemorivagus |  |  |  | *Passalites nemorivagus* |  |
|  | Saimiri boliviensis |  |  |  | *Saimiri boliviensis* |  |
|  | Tapirus terrestris |  |  |  | *Sapajus apella* |  |
|  |  |  |  |  | *Tapirus terrestris* |  |
| Trail seg A1 | Ateles chamek | 10 | *Saimiri boliviensis* | 1 | *Ateles chamek* | 12 |
|  | Cebuella niveiventris |  |  |  | *Cebuella niveiventris* |  |
|  | Dicotyles tajacu |  |  |  | *Cebus cuscinus* |  |
|  | Leontocebus weddelli |  |  |  | *Dicotyles tajacu* |  |
|  | Nasua nasua |  |  |  | *Leontocebus weddelli* |  |
|  | Passalites nemorivagus |  |  |  | *Nasua nasua* |  |
|  | Puma concolor |  |  |  | *Passalites nemorivagus* |  |
|  | Saimiri boliviensis |  |  |  | *Puma concolor* |  |
|  | Tapirus terrestris |  |  |  | *Saimiri boliviensis* |  |
|  | Tayassu pecari |  |  |  | *Sapajus apella* |  |
|  |  |  |  |  | *Tapirus terrestris* |  |
|  |  |  |  |  | *Tayassu pecari* |  |
| Trail seg. M2 | Alouatta seniculus | 17 | *Alouatta seniculus* | 7 | *Alouatta seniculus* | 16 |
|  | Aotus nigriceps |  | *Ateles chamek* |  | *Aotus nigriceps* |  |
|  | Artibeus planirostris |  | *Cebuella niveiventris* |  | *Artibeus planirostris* |  |
|  | Ateles chamek |  | *Ctenophryne geayi* |  | *Ateles chamek* |  |
|  | Bassaricyon alleni |  | *Nasua nasua* |  | *Bassaricyon alleni* |  |
|  | Cebuella niveiventris |  | *Potos flavus* |  | *Cebuella niveiventris* |  |
|  | Coendou sp. |  | *Tapirus terrestris* |  | *Coendou sp.* |  |
|  | Dicotyles tajacu |  |  |  | *Mesomys hispidus* |  |
|  | Mesomys hispidus |  |  |  | *Nasua nasua* |  |
|  | Nasua nasua |  |  |  | *Oligoryzomys sp.* |  |
|  | Oligoryzomys sp. |  |  |  | *Potos flavus* |  |
|  | Potos flavus |  |  |  | *Puma concolor* |  |
|  | Puma concolor |  |  |  | *Saimiri boliviensis* |  |
|  | Saimiri boliviensis |  |  |  | *Sapajus apella* |  |
|  | Sapajus apella |  |  |  | *Tapirus terrestris* |  |
|  | Tapirus terrestris |  |  |  | *Tayassu pecari* |  |
|  | Tayassu pecari |  |  |  |  |  |
| Trail seg. A2 | Alouatta seniculus | 13 | *Alouatta seniculus* | 4 | *Alouatta seniculus* | 13 |
|  | Aotus nigriceps |  | *Ateles chamek* |  | *Aotus nigriceps* |  |
|  | Ateles chamek |  | *Puma concolor* |  | *Aotus nigriceps* |  |
|  | Cebuella niveiventris |  | *Saimiri boliviensis* |  | *Ateles chamek* |  |
|  | Dicotyles tajacu |  |  |  | *Cebuella niveiventris* |  |
|  | Leptodactylus stenodema |  |  |  | *Dicotyles tajacu* |  |
|  | Mazama americana |  |  |  | *Mazama americana* |  |
|  | Molossus molossus |  |  |  | *Molossus molossus* |  |
|  | Puma concolor |  |  |  | *Puma concolor* |  |
|  | Saimiri boliviensis |  |  |  | *Saimiri boliviensis* |  |
|  | Sapajus apella |  |  |  | *Sapajus apella* |  |
|  | Tapirus terrestris |  |  |  | *Tapirus terrestris* |  |
|  | Uroderma bilobatum |  |  |  | *Uroderma bilobatum* |  |
| Trail seg. M3 | Alouatta seniculus | 10 | *Alouatta seniculus* | 5 | *Alouatta seniculus* | 10 |
|  | Ateles chamek |  | *Ateles chamek* |  | *Ateles chamek* |  |
|  | Cebuella niveiventris |  | *Nasua nasua* |  | *Cebuella niveiventris* |  |
|  | Cebus cuscinus |  | *Saimiri boliviensis* |  | *Cebus cuscinus* |  |
|  | Didelphis marsupialis |  | *Tapirus terrestris* |  | *Didelphis marsupialis* |  |
|  | Mesomys hispidus |  |  |  | *Mesomys hispidus* |  |
|  | Nasua nasua |  |  |  | *Nasua nasua* |  |
|  | Oligoryzomys sp. |  |  |  | *Oligoryzomys sp.* |  |
|  | Saimiri boliviensis |  |  |  | *Saimiri boliviensis* |  |
|  | Tapirus terrestris |  |  |  | *Tapirus terrestris* |  |
| Trail seg. A3 | Alouatta seniculus | 12 | *Alouatta seniculus* | 5 | *Alouatta seniculus* | 12 |
|  | Ateles chamek |  | *Ateles chamek* |  | *Ateles chamek* |  |
|  | Cebuella niveiventris |  | *Nasua nasua* |  | *Cebuella niveiventris* |  |
|  | Cebus cuscinus |  | *Saimiri boliviensis* |  | *Cebus cuscinus* |  |
|  | Dicotyles tajacu |  | *Tapirus terrestris* |  | *Dicotyles tajacu* |  |
|  | Eira barbara |  |  |  | *Eira barbara* |  |
|  | Mazama americana |  |  |  | *Mazama americana* |  |
|  | Nasua nasua |  |  |  | *Nasua nasua* |  |
|  | Saimiri boliviensis |  |  |  | *Saimiri boliviensis* |  |
|  | Sapajus apella |  |  |  | *Sapajus apella* |  |
|  | Scinax ictericus |  |  |  | *Scinax ictericus* |  |
|  | Tapirus terrestris |  |  |  | *Tapirus terrestris* |  |
| **Mean # species per trail segment** |  | **12** |  | **4** |  | **12.33** |
| Day 1 | Ateles chamek | 14 | Na | Na | *Ateles chamek* | 16 |
|  | Bassaricyon alleni |  |  |  | *Bassaricyon alleni* |  |
|  | Cebuella niveiventris |  |  |  | *Cebuella niveiventris* |  |
|  | Cuniculus paca |  |  |  | *Cebus cuscinus* |  |
|  | Dasypus sp. |  |  |  | *Cuniculus paca* |  |
|  | Dicotyles tajacu |  |  |  | *Dasypus sp.* |  |
|  | Leontocebus weddelli |  |  |  | *Dicotyles tajacu* |  |
|  | Nasua nasua |  |  |  | *Leontocebus weddelli* |  |
|  | Oligoryzomys sp. |  |  |  | *Nasua nasua* |  |
|  | Passalites nemorivagus |  |  |  | *Oligoryzomys sp.* |  |
|  | Puma concolor |  |  |  | *Passalites nemorivagus* |  |
|  | Saimiri boliviensis |  |  |  | *Puma concolor* |  |
|  | Tapirus terrestris |  |  |  | *Saimiri boliviensis* |  |
|  | Tayassu pecari |  |  |  | *Sapajus apella* |  |
|  |  |  |  |  | *Tapirus terrestris* |  |
|  |  |  |  |  | *Tayassu pecari* |  |
| Day 2 | *Alouatta seniculus* | 21 | Na | Na | *Alouatta seniculus* | 20 |
|  | *Aotus nigriceps* |  |  |  | *Aotus nigriceps* |  |
|  | *Artibeus planirostris* |  |  |  | *Artibeus planirostris* |  |
|  | *Ateles chamek* |  |  |  | *Ateles chamek* |  |
|  | *Bassaricyon alleni* |  |  |  | *Bassaricyon alleni* |  |
|  | *Cebuella niveiventris* |  |  |  | *Cebuella niveiventris* |  |
|  | *Coendou sp.* |  |  |  | *Coendou sp.* |  |
|  | *Dicotyles tajacu* |  |  |  | *Dicotyles tajacu* |  |
|  | *Leptodactylus stenodema* |  |  |  | *Mazama americana* |  |
|  | *Mazama americana* |  |  |  | *Mesomys hispidus* |  |
|  | *Mesomys hispidus* |  |  |  | *Molossus molossus* |  |
|  | *Molossus molossus* |  |  |  | *Nasua nasua* |  |
|  | *Nasua nasua* |  |  |  | *Oligoryzomys sp.* |  |
|  | *Oligoryzomys sp.* |  |  |  | *Potos flavus* |  |
|  | *Potos flavus* |  |  |  | *Puma concolor* |  |
|  | *Puma concolor* |  |  |  | *Saimiri boliviensis* |  |
|  | *Saimiri boliviensis* |  |  |  | *Sapajus apella* |  |
|  | *Sapajus apella* |  |  |  | *Tapirus terrestris* |  |
|  | *Tapirus terrestris* |  |  |  | *Tayassu pecari* |  |
|  | *Tayassu pecari* |  |  |  | *Uroderma bilobatum* |  |
|  | *Uroderma bilobatum* |  |  |  |  |  |
| Day 3 | *Alouatta seniculus* | 15 | Na | Na | *Alouatta seniculus* | 15 |
|  | *Ateles chamek* |  |  |  | *Ateles chamek* |  |
|  | *Cebuella niveiventris* |  |  |  | *Cebuella niveiventris* |  |
|  | *Cebus cuscinus* |  |  |  | *Cebus cuscinus* |  |
|  | *Dicotyles tajacu* |  |  |  | *Dicotyles tajacu* |  |
|  | *Didelphis marsupialis* |  |  |  | *Didelphis marsupialis* |  |
|  | *Eira barbara* |  |  |  | *Eira barbara* |  |
|  | *Mazama americana* |  |  |  | *Mazama americana* |  |
|  | *Mesomys hispidus* |  |  |  | *Mesomys hispidus* |  |
|  | *Nasua nasua* |  |  |  | *Nasua nasua* |  |
|  | *Oligoryzomys sp.* |  |  |  | *Oligoryzomys sp.* |  |
|  | *Saimiri boliviensis* |  |  |  | *Saimiri boliviensis* |  |
|  | *Sapajus apella* |  |  |  | *Sapajus apella* |  |
|  | *Scinax ictericus* |  |  |  | *Scinax ictericus* |  |
|  | *Tapirus terrestris* |  |  |  | *Tapirus terrestris* |  |
| **Mean # species per Day** |  | **16.67** |  |  |  | **17** |
